# Representation Learning of Genomic Sequence Motifs with Convolutional Neural Networks

**DOI:** 10.1101/362756

**Authors:** Peter K. Koo, Sean R. Eddy

## Abstract

Although convolutional neural networks (CNNs) have been applied to a variety of computational genomics problems, there remains a large gap in our understanding of how they build representations of regulatory genomic sequences. Here we perform systematic experiments on synthetic sequences to reveal how CNN architecture, specifically convolutional filter size and max-pooling, influences the extent that sequence motif representations are learned by first layer filters. We find that CNNs designed to foster hierarchical representation learning of sequence motifs - assembling partial features into whole features in deeper layers - tend to learn *distributed* representations, *i.e*. partial motifs. On the other hand, CNNs that are designed to limit the ability to hierarchically build sequence motif representations in deeper layers tend to learn more interpretable *localist* representations, *i.e*. whole motifs. We then validate that this representation learning principle established from synthetic sequences generalizes to *in vivo* sequences.

## Introduction

Deep convolutional neural networks (CNNs) have recently been applied to predict transcription factor (TF) binding motifs from genomic sequences (Zhou and Troyanskaya, 2015; Quang and Xie, 2016; Kelley *et al*, 2016; Hiranuma *et al*, 2017). Despite the promise that CNNs bring in replacing methods that rely on *k*-mers and position weight matrices (PWMs) (Ghandi *et al*, 2016; Foat *et al*, 2006), there remains a large gap in our understanding of why CNNs perform well.

A convolutional layer of a CNN is comprised of a set of filters, each of which can be thought of as a PWM. Each filter scans across the inputs, and outputs a non-linear similarity score at each position, a so-called feature map. The filters are parameters of the CNN that are trained to detect relevant patterns in the data. Deep CNNs are constructed by feeding the feature maps of a convolutional layer as input to another convolutional layer. This can be repeated to create a network with any depth. CNNs typically employ *max-pooling* after each convolutional layer, which down-sample the feature maps by setting non-overlapping windows with a single maximum score, separately for each filter. Max-pooling enables deeper layers to detect features hierarchically across a larger spatial scale of the input. CNN predictions are then made by feeding the feature map of the final convolutional layer through one or more fully-connected hidden layers followed by a final output layer.

In genomics, it is unclear how CNN architecture influences the representations of sequence motifs learned throughout the network. Previous studies have suggested that first layer convolutional filters learn representations of sequence motifs, while deeper layers learn combinations of these motifs, so-called regulatory grammars (Alipanahi *et al*, 2015; Angermueller *et al*, 2016; Zeng *et al*, 2016; Quang and Xie, 2016; Kelley *et al*, 2016). A common method to validate a trained CNN is to demonstrate that first layer filters have learned biologically meaningful representations, *i.e*. PWM-like representations of sequence motifs (Alipanahi *et al*, 2015; Kelley *et al*, 2016; Quang and Xie, 2016; Angermueller *et al*, 2016; Cuperus *et al*, 2017; Chen *et al*, 2018; Kelley *et al*, 2018; Bretschneider *et al*, 2018; Ben-Bassat *et al*, 2018; Wang *et al*, 2018; Gao *et al*, 2018; Trabelsi *et al*, 2019). The few studies that perform a quantitative motif comparison of the first layer filters against a motif database find that less than 50% have a statistically significant match (Kelley *et al*, 2016; Quang and Xie, 2016). Unmatched filters have been suggested to be either partial representations of known motifs or novel motifs, *i.e*. motifs not included in the database. Visualization of deeper layer filters is challenging, because they represent patterns of feature maps in previous layers, where the spatial positions of activations are obscured by pooling.

Learning whole motif representations by first layer filters is not indicative of a deep CNN’s classification performance. For instance, a deep CNN that employs a small first layer filter, *i.e*. 8 nts (Zhou and Troyanskaya, 2015), which is shorter than many common motifs found *in vivo*, has demonstrated comparable performance as CNNs that employ larger filters, *i.e*. ≥ 19 nts (Quang and Xie, 2016; Kelley *et al*, 2016). In principle, smaller filters that capture partial motif representations can be combined in deeper layers to assemble whole motif representations, thereby allowing the CNN to make accurate predictions. It remains unclear to what extent we should expect first layer filters to learn whole motif representations in the first convolutional layer and how a CNN;s architecture influences this.

Alternative approaches to visualize a CNN;s learned representations can be achieved via attribution methods, such as *in silico* mutagenesis (Alipanahi *et al*, 2015), saliency analysis (Simonyan *et al*, 2013), smoothgrad (Smilkov *et al*, 2017), DeepLift (Shrikumar *et al*, 2017), and SHAP (Lundberg and Lee, 2017). These attribution methods are able to integrate distributed representations throughout the network to arrive at the importance of each nucleotide variant for a given sequence. However, they only consider one sequence at a time and their scores are noisy (Kindermans *et al*, 2017; Adebayo *et al*, 2018), which can result in significant scores for nucleotide variants that are not necessarily biologically relevant. To address these issues, TF-MoDISco splits the attribution scores of each sequence into subsequences called sequlets, clusters and aligns these sequlets, then averages each cluster to provide more interpretable representations learned by the network (Shrikumar *et al*, 2018). The reliability of attribution scores to faithfully recapitulate features learned by a deep CNN remains an open problem (Koo *et al*, 2019).

Interpretability of CNNs can also be improved through design principles that guide optimization methods toward a biologically-meaningful parameter space. Here we demonstrate how architectural choice affects representation learning of genomic sequence motifs. We perform systematic experiments to empirically demonstrate that a CNN;s design, specifically max-pooling and filter size, is indicative of the extent that motif representations are learned in first layer filters. We then demonstrate that the same representation learning principles generalize to *in vivo* sequences. Together, this study enables design of CNNs that intentionally learn interpretable representations in easier to access first layer filters (with a small tradeoff in performance), versus building harder to interpret distributed representations, both of which have their strengths and limitations.

## Results

### Internal representations of motifs depend on architecture

We conjecture that motif representations learned in first layer filters are largely influenced by a CNN’s ability to assemble whole motif representations in deeper layers, which is determined by architectural constraints set by: 1. the convolutional filter size, 2. the stride of the filter, which is the offset between successive applications of the filter (usually set to 1), 3. the max-pool size, and 4. the max-pool stride, which is the offset for each max-pool application. Although these are hyperparameters that may vary on a case-by-case basis, the max-pool stride is commonly set to the max-pool size in genomics, creating non-overlapping max-pool windows.

Despite the complexity of *in vivo* TF binding (Siggers and Gordan, 2013), for the purposes of this paper we make a simplifying assumption that TF binding sites can be represented by a single PWM-like motif pattern. Of course, a PWM-based method would perform well in this over-simplified scenario. However, the scope of this paper is to demonstrate how representations of sequence patterns are learned by a CNN and not a thorough demonstration of a CNN’s ability to learn *in vivo* binding sites of TFs.

Assuming that accurate classification can only be made if the correct motifs are detected, a CNN that learns partial motif representations in the first layer must assemble whole motif representations at some point in deeper layers. To help explain how architecture can influence representation learning in a given layer, we use the concept of a receptive field, which is the sensory space of the data that affects a given neuron’s activity. For the first convolutional layer, each neuron’s receptive field has a size that is equal to the filter size at a particular region of the data. Since there are typically many filters in a convolutional layer, there are many neurons which have a receptive field that share the same spatial region. However, each neuron’s activation is determined by a different filter. Max-pooling combines multiple neurons of a given filter within a specified window size to a single max-pooled neuron, thereby augmenting the size of its receptive field. In doing so, max-pooling obfuscates the exact positioning of the max-activation within each window. Thus the location of the max-activation has spatial invariance within its receptive field with an amount equal to the max-pool size.

Although max-pooling creates spatial uncertainty of the max-activation within a max-pooled neuron’s receptive field, we surmise that neighboring max-pooled neurons of different filters, which share significantly overlapping receptive fields, can help to resolve spatial positioning of an activation. To illustrate, Figure 1A shows a toy example of two convolutional filters, each 7 nts long, which have learned partial motifs: ‘GTG’ and ‘CAC’. An example sequence contains three embedded patterns (highlighted in green): ‘CACGTG’, ‘GTGCAC’, and ‘CACNNNGTG’, where ‘N’ represents any nucleotide with equal probability. The resultant max-pooled, activated convolutional scans for each filter are shown above the sequence with a blue shaded region highlighting the receptive field of select max-pooled neurons. Even though the first convolutional layer filters have learned partial motifs, the second convolutional layer filters can still resolve each of the three embedded patterns by employing filters of length 3. Of course situations may arise where the three second convolutional layer filters are unable to fully resolve the embedded patterns with fidelity. For instance, ‘CACNGTG’ could be activated by the same filter for ‘CACGTG’. A CNN can circumvent these ambiguous situations by either learning more information about each pattern within each filter or by dedicating additional filters to help discriminate the ambiguous patterns.

**Figure 1:**
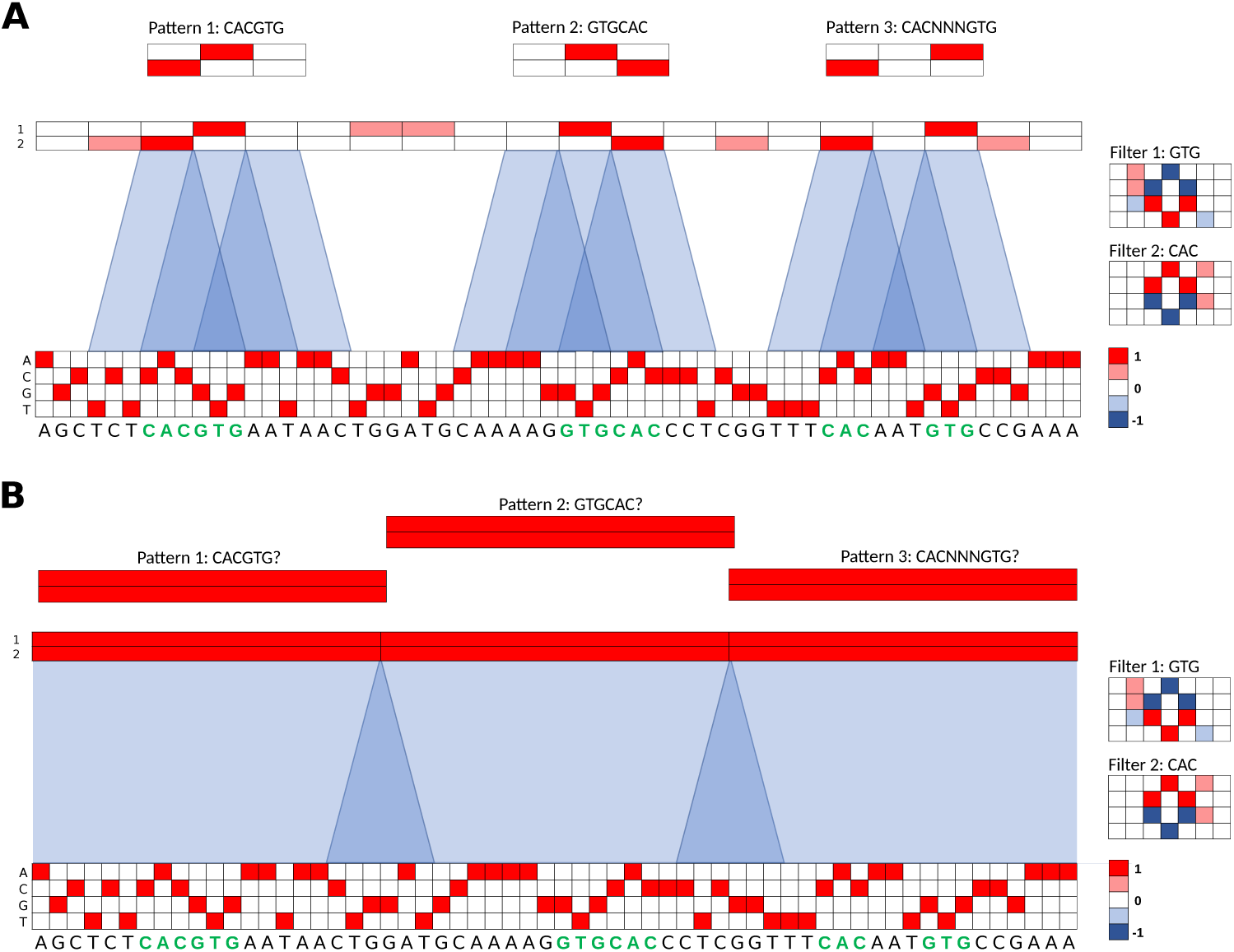
Toy model for representation learning of sequence motifs. (A,B) An example 60 nt one-hot encoded sequence contains 3 patterns (shown in green): CACGTG, GTGCAC, and CACNNNGTG. Two filters, each of length 7 (7 columns and 4 rows, one for each nucleotide), are shown to the right. A partial motif representation has been captured by each filter: GTG for filter 1 (Top) and CAC for filter 2 (Bottom). The max-pooled feature maps are shown above the sequence. The feature maps have the same size as the sequence by adding 3 zero-padding units to each end of the sequence prior to convolution (not shown in diagram). (A) Shows the feature maps when employing a small max-pooling size of 3, which creates overlapping receptive fields, highlighted in blue. 3 second layer convolutional filters, shown above, demonstrate a feature map pattern that can resolve each embedded sequence pattern. (B) Shows the feature maps when employing a larger pooling size of 20 using the same filters as (A). The larger receptive fields have a large spatial uncertainty along with a small overlap in receptive fields from neighboring neurons. Each of the 3 second layer convolutional filters, shown above, is unable to find a unique feature map pattern that can resolve any embedded sequence pattern.

It follows that by creating a situation where partial motif representations cannot be assembled into whole motifs in deeper layers, learning whole motifs by first layer filters becomes obligatory for accurate classification. One method to limit the information flow through a CNN is by employing large max-pool sizes relative to the filter size. The max-pooled neurons then have large receptive fields with a large spatial uncertainty and only a small overlap in receptive fields with neighboring neurons of different filters. A deeper layer would be unable to resolve the spatial ordering of partial motifs to assemble whole motifs with fidelity. To exemplify, figure 1B shows a toy example of a CNN that employs a larger pool size of 20. Importantly, there are large spatial regions within a receptive field for which a neighboring neuron cannot help to resolve due to a lack of overlap in receptive fields. As a result, deeper convolutional layer filters which are dedicated to each pattern would yield the same signature, unable to resolve any of the three patterns.

More technically, the extent of motif information that each filter learns is guided by the gradients of the objective function, which serves as a measure of the classification error. Assuming accurate classification can only be achieved upon discriminating the underlying motifs of each class, once whole motifs for each class are learned, then the objective function is minimized and the training gradients go to zero. If a CNN can build whole motifs from partial motifs in deeper layers, then there is no more incentive to learn additional information to build upon the partial motif representations already learned. As a result, the first layer filters will maintain a *distributed* representations of motifs (Hinton *et al*, 1986). On the other hand, if architectural constraints limit the ability to build whole motifs from partial motifs in deeper layers, then accurate predictions cannot be made. Hence, gradients will persist because the objective function is not yet minimized, encouraging first layer filters to learn whole motifs, also known as a *localist* representation of motifs (Hinton *et al*, 1986). Once the first layer filters have learned sufficient information of whole motifs to discriminate each class, then the objective function can be minimized, signaling the end of training.

### Max-pooling influences ability to build hierarchical motif representations

To test this idea, we created a synthetic dataset that enforces our simplifying assumption that the only important pattern for a given TF binding event is the presence of a PWM-like motif in a sequence. Briefly, synthetic sequences, each 200 nts long, were implanted with 1 to 5 known TF motifs, randomly selected with replacement from a pool of 12 transcription factor motifs embedded in random DNA (see Methods for details). The motifs were manually selected from the JASPAR database to represent a diverse, non-redundant set. The goal of this computational task is to simultaneously make 12 binary predictions for the presence or absence of each transcription factor motif in the sequence. Since we have ground truth for all of the relevant TF motifs and where they are embedded in each sequence, we can test the efficacy of the representations learned by a trained CNN. We note that the ground truth is only from embedded motifs and not from motifs that occasionally arise by chance; the latter effectively creates false negative labels in this dataset.

A CNN that employs at least two convolutional layers is required to test our hypotheses of representation learning. We constructed a CNN with 3 hidden layers: two convolutional layers, each followed by max-pooling, and a fully-connected hidden layer. Specifically, our CNN takes as input one-hot encoded sequences, processes them with the hidden layers, and outputs a prediction for the binding probability for each of the 12 classes. The number of filters in each convolutional layer, the number of units in the fully-connected hidden layer, and the dropout probabilities are fixed (see Methods). The filter sizes, the max-pool window sizes, and the max-pool strides are the hyperparameters that can be varied. For a given hyperparameter setting, we trained the CNN as a multi-class logistic regression (see Methods for training details). All reported metrics are strictly drawn from the held-out test set using model parameters that yielded the best performance on the validation set.

To explore how spatial uncertainty within receptive fields set by max-pooling influences the representations learned by first layer filters, we systematically altered the max-pool sizes while keeping all other hyperparameters fixed, including a first and second layer filter size of 19 and 5, respectively. To minimize the influence of architecture on classification performance, we coupled the max-pool sizes between the first and second layer, such that their products are equal, which makes the inputs into the fully-connected hidden layer the same size across all CNNs. The max-pool sizes we employed are (first layer, second layer): (1,100), (2, 50), (4, 25), (10, 10), (25, 4), (50, 2), and (100,1). For brevity, we denote each CNN with only the first max-pool window size, e.g. CNN-2 for (2, 50).

We first verified that the performance of each model is similar as measured by the average area under the receiver-operator-characteristic (AU-ROC) curve across the 12 classes (Table 1), which is in the range of previously reported values for a similar task using experimental ChIP-seq data (Zhou and Troyanskaya, 2015; Quang and Xie, 2016). Next, we converted each filter to a sequence logo to visually compare the motif represenations learned by the first layer filters across the different architectures (see Methods). As expected, we found CNNs that employ large max-pool sizes (≥10) learn representations that qualitatively resemble the ground truth motifs (Fig. 2). On the other hand, CNNs that employ a small max-pool size (≤4) do not seem to qualitatively capture any ground truth motif in its entirety, perhaps learning, at best, parts of a motif.

**Table 1:** Performance on the synthetic dataset. The table shows the average area under the receiver-operator-characteristic curve (AU-ROC) across the 12 TF classes, percentage of matches between the 30 first convolutional layer filters and the entire JASPAR vertebrates database (JASPAR), and the percentage of filters that match to any ground truth TF motif (Relevant) for different CNNs. Motif matches were determined by the Tomtom motif comparison search tool using an *E*-value cutoff of 0.1. The Average AU-ROC error represents the standard deviation of the AU-ROC across the 12 classes.

| Model | Average<br>AU-ROC | % Motif match<br>(JASPAR) | % Motif match<br>(Relevant) |
| --- | --- | --- | --- |
| CNN-1 | 0.972±0.001 | 0.240±0.083 | 0.007±0.013 |
| CNN-2 | 0.966±0.000 | 0.240±0.071 | 0.007±0.013 |
| CNN-4 | 0.955±0.002 | 0.453±0.131 | 0.127±0.080 |
| CNN-10 | 0.964±0.007 | 0.987±0.016 | 0.973±0.033 |
| CNN-25 | 0.973±0.001 | 0.987±0.016 | 0.980±0.027 |
| CNN-50 | 0.961±0.011 | 0.933±0.037 | 0.920±0.045 |
| CNN-100 | 0.958±0.012 | 0.887±0.034 | 0.880±0.034 |
| CNN<sub>9</sub>-4 | 0.954±0.002 | 0.260±0.039 | 0.033±0.030 |
| CNN<sub>9</sub>-25 | 0.958±0.008 | 0.993±0.013 | 0.980±0.016 |
| CNN<sub>3</sub>-50 | 0.648±0.008 | 0.160±0.049 | 0.000±0.000 |
| CNN<sub>3</sub>-2 | 0.968±0.001 | 0.233±0.067 | 0.000±0.000 |
| CNN-50-2 | 0.921±0.012 | 0.913±0.050 | 0.893±0.044 |
| CNN<sub>19-1</sub>-2 | 0.969 ±0.002 | 0.867±0.056 | 0.747±0.096 |
| CNN-25 (60) | 0.972±0.001 | 0.973±0.013 | 0.960±0.023 |
| CNN-25 (90) | 0.968±0.001 | 0.940±0.023 | 0.909±0.028 |
| CNN-25 (120) | 0.963±0.001 | 0.933±0.015 | 0.887±0.025 |

**Figure 2:**
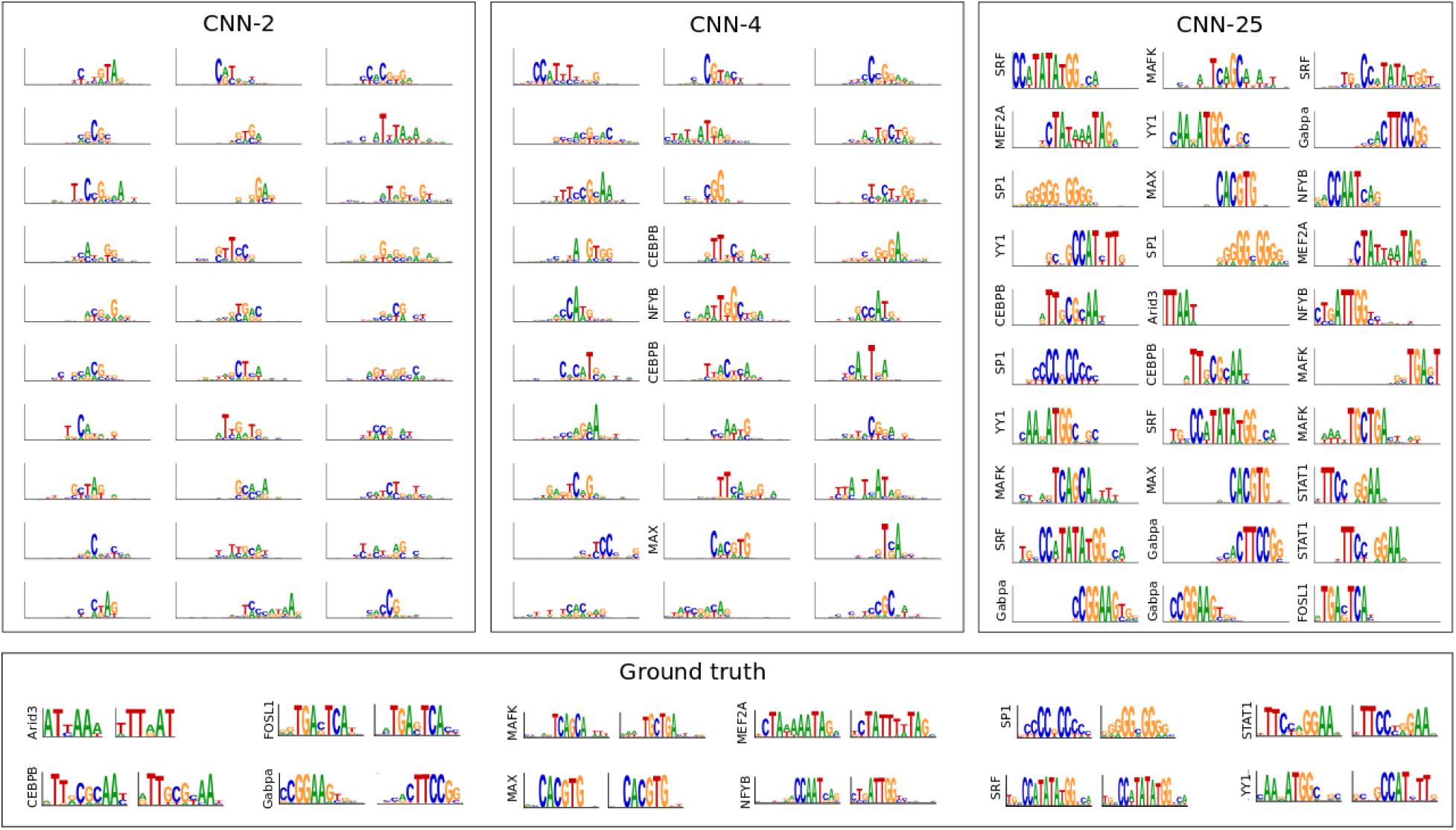
Comparison of first layer filters for CNNs with a different max-pool size. Sequence logos for first convolutional layer filters are shown for CNN-2 (Left), CNN-4 (Middle), and CNN-25 (Right). The sequence logo of the ground truth motifs and its reverse complement for each transcription factor is shown at the bottom. The y-axis label on select filters represents a statistically significant match to a ground truth motif.

To quantify the number of filters that have learned motifs, we employed the Tomtom motif comparison search tool (Gupta *et al*, 2007) to compare the similarity of each filter against all motifs in the JASPAR 2016 vertebrate database (Mathelier *et al*, 2016) using an E-value cutoff of 0.1. In agreement with our qualitative observation, we found that the fraction of filters that match known motifs for CNNs that employ a small max-pool size (≤4) yield, at best, 0.453 ±0.131 (error is the standard deviation of the mean across 5 independent trials). Of these, only a small fraction match any ground truth motif (0.127 ±0.080). The large discrepancy between the number of filters that match any motif in the JASPAR database versus the number that match ground truth motifs demonstrates that motif comparison is not a reliable tool to identify whether a CNN learns *relevant* motifs. In this study, we know what the ground truth is and hence can reliably calculate this quantity.

In contrast, CNNs that employ a large max-pool size yield, at worst, a match fraction of 0.880 ±0.034 to ground truth motifs, with CNN-25 yielding the highest fraction of 0.980 ±0.027 (Table 1). These results support our hypothesis that limiting the ability to build hierarchical representations with a spatial information bottleneck created by max-pooling encourages first layer filters to learn whole motif representations.

Even when a CNN is designed to learn whole motif representations, not all filters learn motifs, and the ones that do may learn slightly different versions of the same motif. This variation does not necessarily represent real differences in the underlying data. Also, the redundancy – number of filters dedicated to a motif – is not necessarily a reliable measure of its importance. The variability in the number of filters dedicated to a motif in this dataset, which do not have any class imbalance, suggests that this observed difference is more likely due to the difficulty of finding that motif from random initialization, not the importance of the motif.

### Sensitivity of motif representations to the number of filters

To test the sensitivity of motif representations to the number of first layer filters for CNN-25, we systematically increased the number of first layer filters in the first convolutional layer of CNN-25 from 30 to 60, 90, and 120. Upon training each of these models, we found that the number of filters that match relevant motifs is 0.980 ±0.027, 0.960 ±0.023, 0.909 ±0.028, 0.887 ±0.025, respectively (Table 1). Overparameterizing the number of first layer filters results more filters that do not learn any motif representations. We attribute this to the ease of finding relevant motifs from random initializations. As training begins, each filter is guided by gradient descent to find relevant patterns that help to minimize the objective function. When the network is overparameterized, there are many filters randomly initialized, some of which may be in a better position to learn relevant motifs. As these filters learn relevant motifs first, the objective function becomes minimized, leaving the remainder of the filters without any gradients to finish learning a relevant motif.

### Motif representations are not very sensitive to 1st layer filter size

Due to the common misconception that first convolutional layer filters learn motifs (Alipanahi *et al*, 2015; Kelley *et al*, 2016; Quang and Xie, 2016; Angermueller *et al*, 2016; Cuperus *et al*, 2017; Chen *et al*, 2018; Kelley *et al*, 2018; Bretschneider *et al*, 2018; Ben-Bassat *et al*, 2018; Wang *et al*, 2018; Gao *et al*, 2018; Trabelsi *et al*, 2019), deep learning practitioners continue to employ CNN architectures with large first layer filters with the intent of capturing motif patterns in their entirety. However, we have shown that employing a large filter does not necessarily lead to whole motif representations. To test the sensitivity of filter size to representation learning, we created two new CNNs that employ a first layer filter size of 9 (CNN _9_), in contrast to a filter size of 19 which was previously used, with max-pool combinations of 4 and 25, *i.e*. CNN_9_-4 and CNN_9_-25. Since the combination of a filter size of 9 with a max-pool size of 4 creates overlapping receptive fields with a small spatial uncertainty, we expect that this architecture setting will lead to partial motif representations. On the other hand, the filter size of 9 is insufficient to resolve spatial positions when employing a max-pool size of 25. Hence, we predict that this architecture setting will yield whole motif representations. As expected, CNN_9_-25 learns representations that qualitatively better reflect the ground truth motifs compared to CNN_9_-4 (Fig. 3, A-B), albeit only learning partial motif representations of larger motifs, *i.e*. MEF2A, SRF, STAT1, CEBPB, but in a more visually identifiable way compared to CNN _9_-4. By quantifying the percentage of filters that statistically match ground truth motifs, CNN _9_-25 yields an 93% match compared to CNN_9_-4 which yield no matches (Table 1). Performance of CNN_9_-25 is lower than CNN-25, likely due to learning more complete (wider) motifs. Due to random initialization, wide first layer filters are preferable to capture the whole motif.

**Figure 3:**
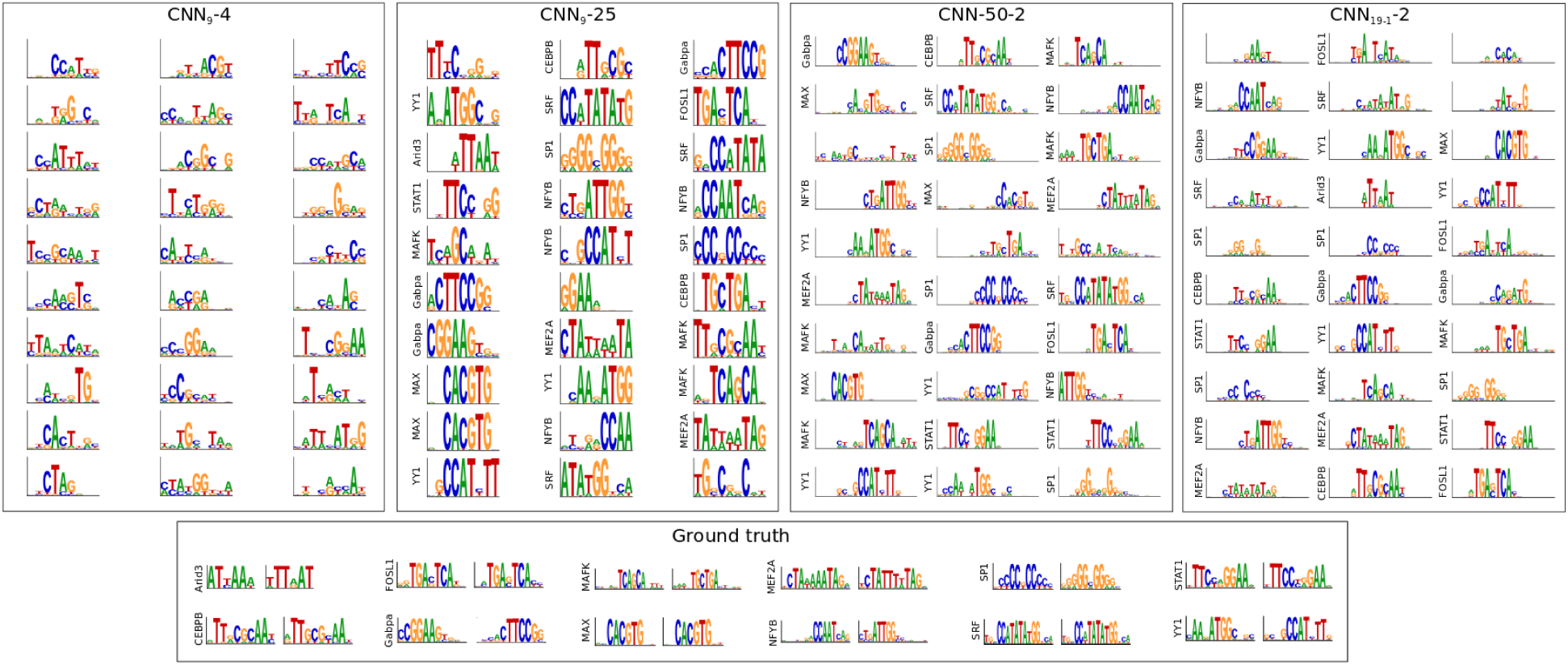
Representations learned by first layer filters for alternative CNN architectures. Sequence logos for first convolutional layer filters are shown for (from left to right): CNN _9_-4, CNN_9_-25, CNN-50-2, and CNN_19-1_-2. The sequence logo of the ground truth motifs and its reverse complement for each transcription factor is shown at the bottom. The y-axis label on select filters represent a statistically significant match to a ground truth motif.

As a control, we created an extreme CNN architecture that would likely not be used in practice if one performed hyperparameter optimization: CNN_3_-2 and CNN_3_-50, which has a first layer filter size of 3 and max-pool size combinations of 2 and 50, respectively. Even though the filter size is smaller than many of the embedded motifs, we expect that CNN_3_-2 will still be able to assemble whole motifs to some extent in deeper layers, because it employs small max-pooling. On the other hand, since CNN _3_-50 has only one chance to learn whole motifs, we expect that the small filter size will limit its ability to discriminate the embedded motifs for each class, leading to a poor classification performance. Indeed, CNN _3_-50 yields a mean AU-ROC of 0.652±0.060 across the 12 classes, compared to CNN_3_-2 which yields 0.968±0.039.

### Motif representations are affected by the ability to assemble whole motifs in deeper layers

One aspect of max-pooling that we did not consider in our toy model is the max-pool stride, which is typically set to the max-pool size. Employing a large max-pool size with a small max-pool stride can create a situation where the receptive field of max-pooled neurons overlap significantly, which should improve the spatial resolution of partial motifs. However, a deeper convolutional filter would still be unable to assemble whole motifs, because each receptive field has a large spatial uncertainty, which provides the same activation pattern irrespective of whether partial motifs are close together or very distant. Hence, we expect this CNN to learn whole motif representations.

To test this, we created a new CNN which employs a large max-pool size of 50 with a max-pool stride of 2 (CNN-50-2). Consequently, the length of the feature maps after the first convolutional layer are half of the input sequence, which is the same shape as the feature maps of CNN-2, which employs a max-pool size of 2 with a stride of 2. Similar to CNN-2, CNN-50-2 employs a max-pool size and stride of 50 after the second convolutional layer, which ensures that both networks have the same number of parameters. As expected, CNN-50-2 learns whole motif representations with 90% of its filters matching ground truth motifs in the synthetic dataset (Table 1). Moreover, the motifs that are learned by CNN-50-2 qualitatively better resemble whole motif representations (Fig. 3) compared to CNN-2 (Fig. 2). Together, this result further supports that architecture, specifically the ability to assemble whole motifs in deeper layers, plays a major role in how CNNs learn genomic representations in a given layer.

Another factor that can affect the ability to assemble whole motifs in deeper layers is the size of second convolutional layer filters. A small filter size can make it challenging to assemble whole motifs, even if the max-pool size and stride are small. To test this, we modified the second convolutional filter size of CNN-2 from a size of 5 to 1 (CNN_19-1_-2). As expected, the first layer filters learn representations of whole motifs (87% match to ground truth motifs) even though CNN_19-1_-2 employs a small max-pool size of 2 (Fig. 3 and Table 1). This demonstrates that motif representations learned in a given layer are not only affected by max-pooling but also the ability of deeper layers to hierarchically build whole motif representations.

### Distributed representations build whole motif representations in deeper layers

The high classification performance of each CNN suggests that it must have learned whole motif representations at some point. Thus, CNNs, whose first layer filters do not match any relevant motifs, must be assembling whole motif representations in deeper layers. Alignment-based visualization of filters is not straightforward for filters in deeper layers, because max-pooling obfuscates the exact positions of activations. However, CNN-1 does not employ any max-pooling after the first convolutional layer, and hence can use the same alignment-based visualization approach for second layer filters. The fraction that CNN-1’s 128 second layer filters match any motif in the JASPAR database and ground truth motifs is 0.900 ±0.024 and 0.847 ±0.021, respectively (Fig. S1). On the other hand, CNN-1’s first layer filters yielded 0.240 ±0.083 and 0.007 ±0.013. This demonstrates that second layer filters indeed are fully capable of assembling partial motif representations into whole motif representations.

To test whether second layer filters always build whole motif representations when first layer filters learn distributed representations, we designed another CNN with 3 convolutional layers (CNN-1-1-100). CNN-1-1-100 augments CNN-1 with an additional convolutional layer (128 filters of size 5 and no max-pooling) after the first convolutional layer. The first max-pooling is applied after the 3rd convolutional layer with a pool size of 100, and hence the third convolutional layer now sets the bottleneck for spatial information. CNN-1-1-100’s match fraction to the JASPAR database and to ground truth motifs are: 0.147 ±0.045 and 0.0 ±0.0 for first layer filters; 0.192 ±0.022 and 0.006 ±0.006 for second layer filters; and 0.927 ±0.020 and 0.89, ±0.030 for third layer filters. Hence, the layer which sets the large spatial information bottleneck is where the majority of whole motif representations are learned. This empirical result supports that this representation learning principle also generalizes to deeper layers.

### Generalization to *in vivo* sequences

To test whether the same representation learning principles generalize to *in vivo* sequences, we modified the DeepSea dataset (Zhou and Troyanskaya, 2015) to include only *in vivo* sequences that have a peak called for at least one of 12 ChIP-seq experiments, each of which correspond to a TF in the synthetic dataset (see Table **??**). The truncated-DeepSea dataset is similar to the synthetic dataset, except that the input sequences now have a size of 1,000 nt in contrast to the 200 nt sequences in the synthetic dataset.

We trained each CNN on the *in vivo* dataset following the same protocol as the synthetic dataset. Based on a simplifying assumption that *in vivo* TF binding sites can be captured by a single PWM-like motif patterns, we compare the first layer filters of each CNN. Similar to the results from the synthetic dataset, a qualitative comparison of the first layer filters of different CNNs show that employing a larger pool size yield representations that better reflect whole motifs (Fig. 4). Since *in vivo* sequences are not necessarily accompanied with ground truth, we were unable to reliably quantify the percentage of filters that learn ground truth motifs. Instead, we quantified the fraction of filters that match the known motifs associated with the TF in each ChIP-seq experiment, which we refer to as *relevant* motifs. As expected, first layer filters for CNNs with larger pool sizes better capture relevant motif representations (see Table 2). On the other hand, CNN-1, which does not employ any max-pooling, yielded a match fraction of 0.227 ±0.068 to any motif in the JASPAR database and 0.020 ±0.027 to relevant motifs. In agreement with results on synthetic data, CNN-1’s second layer filters yielded a significantly higher match fraction of 0.845 ±0.012 and 0.625 ±0.030 to any motif in the JASPAR database and to relevant motifs, respectively (see Fig. S2). This further supports that deeper layers are able to assemble partial representations into deeper layers.

**Table 2:** Performance of deep learning models on the *in vivo* dataset. The table shows the average area under the receiver-operator-characteristic curve (AU-ROC) and the average area under the precision recall curve (AU-PR) across the 12 TF classes, percentage of matches between the 30 first convolutional layer filters and the entire JASPAR vertebrates database (JASPAR), and the percentage of filters that match to any ground truth TF motif (Relevant) for different CNNs. Motif matches were determined by the Tomtom motif comparison search tool using an *E*-value cutoff of 0.1. The average AU-ROC error and the average AU-PR error represents the standard deviation across the 12 classes.

| Model | Average<br>AU-ROC | Average<br>AU-PR | Motif match<br>(JASPAR) | Motif match<br>(Relevant) |
| --- | --- | --- | --- | --- |
| CNN-1 | 0.918±0.001 | 0.626±0.000 | 0.227±0.068 | 0.020±0.027 |
| CNN-2 | 0.911±0.003 | 0.609±0.006 | 0.333±0.067 | 0.107±0.057 |
| CNN-4 | 0.907±0.002 | 0.601±0.005 | 0.753±0.045 | 0.507±0.039 |
| CNN-10 | 0.903±0.006 | 0.583±0.020 | 0.920±0.045 | 0.753±0.034 |
| CNN-25 | 0.903±0.003 | 0.580±0.009 | 0.933±0.030 | 0.747±0.040 |
| CNN-50 | 0.903±0.003 | 0.582±0.009 | 0.913±0.034 | 0.733±0.063 |
| CNN-100 | 0.889±0.004 | 0.554±0.009 | 0.873±0.039 | 0.740±0.057 |
| CNN<sub>9</sub>-4 | 0.906±0.002 | 0.592±0.005 | 0.473±0.061 | 0.187±0.045 |
| CNN<sub>9</sub>-25 | 0.891±0.002 | 0.550±0.008 | 0.880±0.062 | 0.660±0.025 |
| CNN-50-2 | 0.892±0.004 | 0.556±0.010 | 0.853±0.091 | 0.653±0.016 |
| CNN-25 (60) | 0.916±0.003 | 0.621±0.011 | 0.973±0.013 | 0.960±0.023 |
| CNN-25 (90) | 0.919±0.001 | 0.628±0.005 | 0.940±0.023 | 0.909±0.028 |
| CNN-25 (120) | 0.920±0.002 | 0.637±0.005 | 0.933±0.015 | 0.887±0.025 |

**Figure 4:**
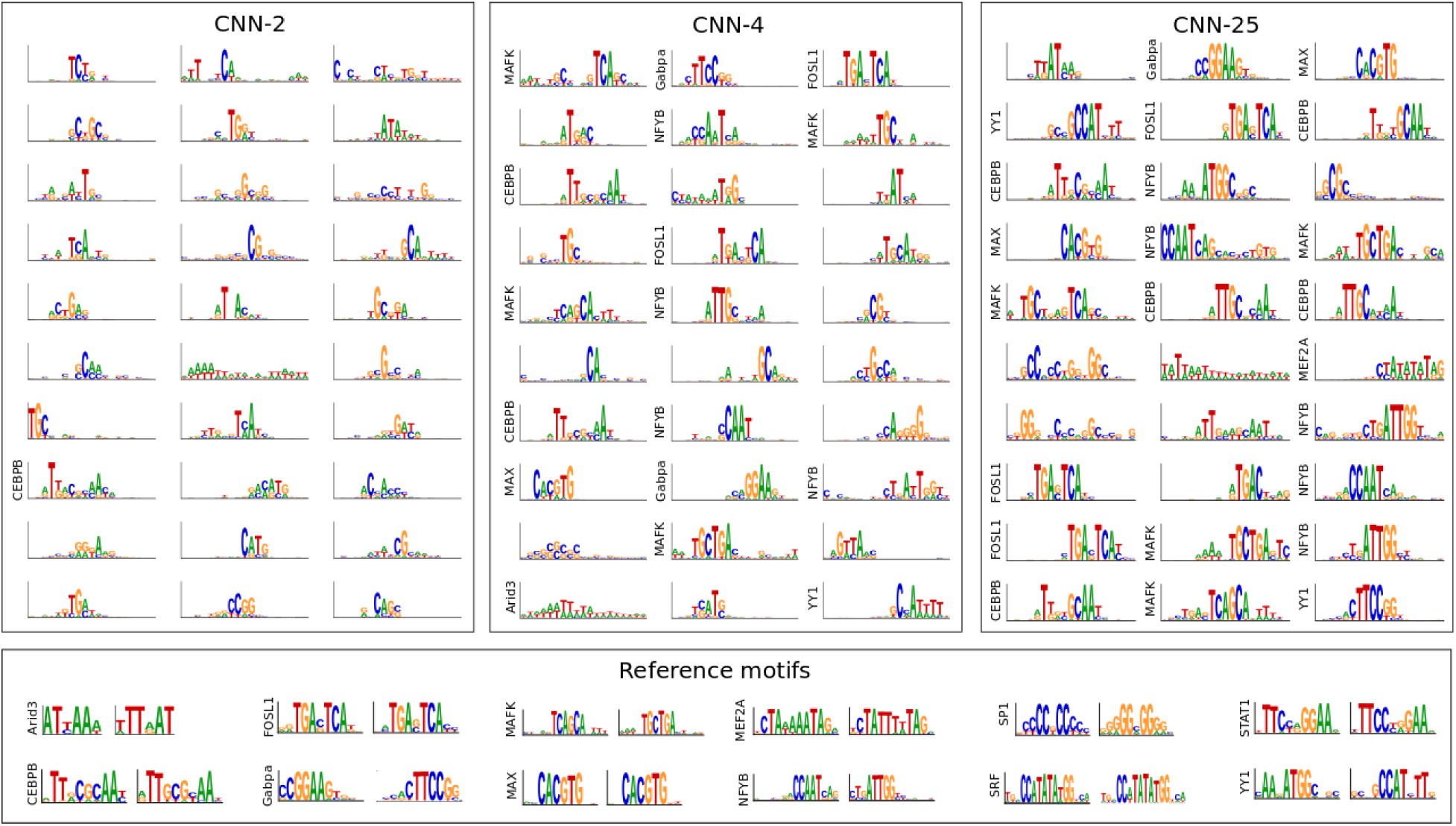
Comparison of the first layer filters for CNNs trained on *in vivo* sequences. Sequence logos for first convolutional layer filters are shown for CNN-2 (Left), CNN-4 (Middle), and CNN-25 (Right). The sequence logos of reference motifs and their reverse complements for each transcription factor from the JASPAR database is shown at the bottom. The y-axis label on select filters represents a Tomtom match to a reference motif.

CNN architectures that are designed to learn distributed representations perform slightly better than the CNN architectures that learn localist representations for *in vivo* sequences. We attribute this performance discrepancy to a CNN’s ability to build a larger repertoire of sequence patterns from distributed representations, which is more important for *in vivo* sequences which have a richer array of biologically important sequence patterns (Siggers and Gordan, 2013). For CNNs that learn localist representations, the number of filters in the first layer sets a hard constraint on the number of sequence patterns that can be detected. To test whether CNN-25’s performance can improve if we enable it to learn more patterns, we performed an experiment where we systematically increased the number of its first layer filters from 30 to 60, 90, and 120. Indeed the performance steadily improves as the number of filters increases (Table 2). Interestingly, the number of filters that match relevant motifs also increases, which suggests that many filters are learning representations of the same motif. The larger number of filters enables diverse initializations which help to learn harder to find motifs - such as ARID3A, SP1, and STAT1 – all of which CNN-25 struggled to detect with 30 filters, but were captured by CNN-1’s second layer filters and CNN-25 when employing 120 filters.

## Discussion

Here, we reveal principles of how architecture design influences representation learning of sequence motifs by exploring different CNN architectures on a synthetic dataset with a known ground truth. Typical deep CNN architectures in genomics employ large filters and small max-pool sizes, which we would expect to learn *distributed* representations of sequence motifs. Hence, visualization of first convolutional layer filters is not as meaningful for these CNNs. However, we showed that *localist* representations, *i.e*. whole motifs, can be learned by constraining the architecture such that the ability of deeper layers cannot reliably assemble hierarchical representations of motifs. Although this study focuses on first layer representations, we believe that the same principles hold for deeper layers in the network. While we only explored the role of architecture in this study, we note that there may be other factors that contribute to the quality of the learned representations, including regularization and optimization algorithms.

While localist representations provide a simple and direct way to visualize interpretable representations learned by the network, building distributed representations may be more beneficial in more complicated tasks, such as *in vivo* TF binding, because a wider array of representations can be combinatorically constructed from partial representations. There becomes less dependence on convolutional filter lengths and numbers of filters as long as there exist deeper layers that can build representations hierarchically. In contrast, building localist representations enforces harder constraints set by the numbers of filters and the filter lengths, limiting the amount of representations and their sizes that can be learned. Thus, the number of first layer filters and the first layer filter size must be set to sufficiently large values to be able to capture the relevant motifs. When the main features in the dataset are simple, such as whether or not a PWM-like motif is present, then CNN architectures that learn localist representations achieve an easier to interpret model that can perform competitively. Nevertheless, distributed representations may be better suited to address more complicated sequence patterns which cannot be easily captured by PWM-like first layer filters.

There are many caveats to interpreting a CNN by just visualizing first layer filters. We showed that comparing filters to a motif database is not a reliable method to assess the quality of motif representations learned by CNNs. Even if filters learn whole motifs, this approach does not inform how the CNN combines these features to make predictions. If the computational task is multi-task classification, there is no correspondence between features and their respective class. Nevertheless, to interpret current attribution methods which are noisy, clustering methods are required (Shrikumar *et al*, 2018), which ultimately result in representations that also suffer from these same issues. We also note that filter visualization does not specify their importance. Some filters that learn uninterpretable representations may not be given much weight by the CNN. Quantifying this is an important problem that remains a direction of future research. Even without a full understanding of what a CNN is learning, identification of relevant motifs can still be informative to generate experimentally testable hypotheses.

In genomics, most CNN designs are inspired by computer vision. However, there are major differences in what features are important between these data modalities. For instance, in natural images, the lower-level features are less informative edges and textures, while higher layer features capture more important shapes of objects. In genomics, the lower-level features are major features of interest, with higher-level features potentially capturing interactions of these lower-level features, so-called regulatory codes. Designing CNNs that allow easy to access first layer representations can provide a straightforward way to identify the pool of motifs that may be relevant for the classification task. Moreover, endowing CNNs that learn localist representations with plentiful first layer filters can provide sufficient representational power to capture the wide repertoire of relevant low-level biological sequence features. Of course, a CNN can still take into account the interactions of these low-level motif features with deeper layers, which we believe should also follow these same representation learning principles.

## Methods

### Synthetic dataset

The synthetic dataset consists of sequences with known motifs embedded in random DNA sequences to mimic a typical multi-class binary classification task for ChIP-seq datasets. We acquired a pool of 24 PWMs from 12 unique transcription factors (forward and reverse complements) from the JASPAR database (Mathelier *et al*, 2016): Arid a, CEBPB, FOSL1, Gabpa, MEF2A, MAFK, MAX, MEF2A, NFYB, SP1, SRF, STAT1, and YY1. For each sequence, we generated a 200 nt random DNA sequence model with equal probability for each nucleotide. 1 to 5 TF PWMs were randomly chosen with replacement and randomly embedded along the sequence model such that each motif has a buffer of at least 1 nucleotide from other motifs and the ends of the sequence. We generated 25,000 sequence models and simulated a single synthetic sequence from each model. A corresponding label vector of length 12, one for each unique transcription factor, was generated for each sequence with a one representing the presence of a TF’s motif or its reverse complement along the sequence model and zero otherwise. The 25,000 synthetic sequences and their associated labels were then randomly split into a training, validation, and test set according to the fractions 0.7, 0.1, and 0.2, respectively.

### *In vivo* dataset

Sequences which contain ENCODE ChIP-seq peaks were downloaded from the DeepSEA dataset via (Zhou and Troyanskaya, 2015). The human reference genome (GRCh37/hg19) was segmented into non-overlapping 200 nt bins. A vector of binary labels for ChIP-seq peaks and DNase-seq peaks was created for each bin, with a 1 if more than half of the 200 nt bin overlaps with a peak region, and 0 otherwise. Adjacent 200 nt bins were then merged to 1,000 nt lengths and their corresponding labels were also merged. Chromosomes 8 and 9 were excluded from training to test chromatin feature prediction performances, and the rest of the autosomes were used for training and validation. We truncated the DeepSea dataset to include only the sequences which contain at least one of 12 transcription factor labels: Arid3a, CEBPB, FOSL1, Gabpa, MEF2A, MAFK, MAX, MEF2A, NFYB, SP1, SRF, STAT1, and YY1 (See Supplementary Table S1 for ENCODE filenames and class indices from the original DeepSea dataset). 270,382 (92%) sequences comprise the training set and 23,768 (8%) sequences comprise the test set. Each 1000 nt DNA sequence is one-hot encoded into a 4×1000 binary matrix, where rows correspond to A, C, G and T.

### CNN Models

All CNNs take as input a 1-dimensional one-hot-encoded sequence with 4 channels (one for each nucleotide: A, C, G, T), then processes the sequence with two convolutional layers, a fully-connected hidden layer, and a fully-connected output layer with 12 output neurons that have sigmoid activations for binary predictions. Each convolutional layer consists of a 1D cross-correlation operation, which calculates a running sum between convolution filters and the inputs to the layer, followed by batch normalization (Ioffe and Szegedy, 2015), which independently scales the features learned by each convolution filter, and a non-linear activation with a rectified linear unit (ReLU), which replaces negative values with zero.

The first convolutional layer employs 30 filters each with a size of 19 and a stride of 1. The second convolutional layer employs 128 filters each with a size of 5 and a stride of 1. All convolutional layers incorporate zero-padding to achieve the same output length as the inputs. Each convolutional layer is followed by max-pooling with a window size and stride that are equal, unless otherwise stated. The product of the two max-pooling window sizes is equal to 100. Thus, if the first max-pooling layer has a window size of 2, then the second max-pooling window size is 50. This constraint ensures that the number of inputs to the fully-connected hidden layer is the same across all models. The fully-connected hidden layer employs 512 units with ReLU activations.

Dropout (Srivastava *et al*, 2014), a common regularization technique for neural networks, is applied during training after each convolutional layer, with a dropout probability set to 0.1 for convolutional layers and 0.5 for fully-connected hidden layers. During training, we also employed L_2_-regularization with a strength equal to 1e-6. The parameters of each model were initialized according to (He *et al*, 2015), commonly known as He initialization.

All models were trained with mini-batch stochastic gradient descent (mini-batch size of 100 sequences) for 100 epochs, updating the parameters after each mini-batch with Adam updates (Kingma and Ba, 2014), using recommended default parameters with a constant learning rate of 0.0003. Training was performed on a NVIDIA GTX Titan X Pascal graphical processing unit with acceleration provided by cuDNN libraries (Chetlur *et al*, 2014). All reported performance metrics and saliency logos are drawn strictly from the test set using the model parameters which yielded the lowest binary cross-entropy loss on the validation set, a technique known as early stopping.

### Visualizing first layer filters

A first layer filter was visualized by scanning it across every sequence in the test set. Sequences whose max activations was less than a cutoff of 70% of the maximum possible activation achievable for that filter were removed. A subsequence the size of the filter is taken about the max activation for each remaining sequence and assembled into an alignment. Subsequences that are shorter than the filter size, because their max activation is too close to the ends of the sequence, are also disregarded. A position probability matrix is created from the alignment and converted to a sequence logo according to: *I*_*i*_ = log_2_(4)+ Σ_*a*_*p*_*i*_(*n*) log_2_*p*_*i*_(*n*), where *p*_*i*_(*n*) is the probability of nucleotide *n* at position *i*.

### Availability

Python scripts to download and process the datasets and TensorFlow code to build, train, and evaluate the CNNs can be found via https://github.com/p-koo/learning_sequence_motifs.

## Acknowledgements

The authors thank Tim Dunn, Soohyun Cho, and Stefan Paul for helpful feedback on the manuscript.

